# Auditory regulation of hippocampal locomotion circuits by a non-canonical reticular-limbic pathway

**DOI:** 10.1101/2025.03.26.645464

**Authors:** Jessica Winne, George Nascimento, Rafael Pedrosa, Margareth Nogueira, Cristiano S. Simões, Klas Kullander, Katarina E. Leão, Richardson N. Leão

## Abstract

The ability to rapidly detect and respond to unexpected auditory stimuli is critical for adaptive behavior, especially during locomotion. Since movement suppresses auditory cortical activity, it remains unclear how salient auditory information influences locomotor circuits. In this work, using in vivo calcium imaging, electrophysiology, chemo- and optogenetics, we investigate the path that relays loud broadband sounds to the dorsal hippocampus (dHPC) and modulates theta oscillations. We demonstrate that noise accelerates theta frequency and decreases its power, effects mediated by entorhinal cortex (EC) and medial septum (MS) inputs while independent of the primary auditory cortex. Activation of dorsal cochlear nucleus (DCN) neurons projecting to the pontine reticular nucleus (PRN) mimics noise-driven hippocampal responses, supporting a brainstem-limbic auditory processing route. Furthermore, noise selectively modulates CA1 pyramidal neuron and interneuron activity, reflecting diverse circuit dynamics. Finally, loud broadband noise stimulus increased theta coherence between the dHPC and the medial prefrontal cortex (mPFC), enhancing interregional synchronization. These results highlight the mechanisms in which the DCN filters behaviorally relevant sounds promoting acoustic motor integration in the hippocampus during locomotion, without direct influence of the auditory cortex.

## Introduction

The capacity to detect and interpret unexpected auditory stimuli during exploration is vital for survival. Behaviors such as foraging, mating, and navigation often occur in dynamic environments where sensory inputs must be integrated with emotional and motor responses to prioritize adaptive actions [1–3]. However, movement suppresses activity in the auditory cortex (AC), with excitatory neurons in the AC decreasing firing rates during grooming, vocalization, and locomotion, and thereby altering sensory responsiveness [3–6]. Thereby it is not well understood how the brain processes unexpected, potentially threatening acoustic signals to generate rapid and appropriate behavioral responses during locomotion.

Recently, a non-canonical central reticular-limbic auditory pathway, bypassing the auditory cortex, has been shown to be the pathway by which threatening auditory cues contribute to fear conditioning [7,8]. This pathway branches from the cochlear nucleus (CN) through the caudal pontine reticular nucleus (PRN), the pontine central gray (PCG), the medial septum (MS) and reaches the entorhinal cortex (EC) [8]. Neurons along this route selectively respond to high-intensity broadband noise and are crucial for fear conditioning [8,9]. By rapidly relaying auditory information to multimodal integration centers, this pathway supports the prioritization of behaviorally relevant stimuli during locomotion, despite the reduction in auditory cortical activity.

The dorsal hippocampus (dHPC) and entorhinal cortex (EC) are central to spatial navigation and contextual encoding, with neuronal populations such as grid, place, and speed cells coordinating activity to construct cognitive maps of the environment [10–19]. Additionally, the medial septum (MS) integrates spatial and navigational abilities, balancing attention and path integration during navigation and pacing theta oscillations of the dorsal hippocampus (dHPC) and entorhinal cortex (EC) [20–30]. Furthermore, theta oscillations, modulated by EC and MS [23,30–32], are strongly associated with locomotion and are influenced by speed and sensory input factors [33–38] still their integration of non-spatial sensory inputs, such as auditory cues, during active behaviors remains less understood [39–42].

Here we show how loud noise travels through the reticule-limbic pathway and influences dHPC. We employ in vivo calcium imaging, electrophysiology, chemo- and optogenetics to explore, at cellular and circuit level, the neural pathway that potentially frightening sounds use to modulate dHPC output. We discovered that loud noise experienced during locomotion is processed through a non-canonical reticular-limbic auditory pathway, which influences theta oscillations and modulates the activity of specific subgroups of dHPC pyramidal cells and interneurons. Our study offers new insights into integration of unexpected acoustic inputs during adaptive neural processing.

## Results

### Broadband loud noise increases mean theta frequency while decreasing theta and slow gamma power during running

To study how environmental cues such as loud broadband noise modulate hippocampal oscillations, we implanted mice with custom-made intrahippocampal array electrodes in the dHPC and, 10 days later, induced theta oscillations by letting mice run on a treadmill. Electrode placement was routinely confirmed post hoc (Fig. 1A) but also by producing cross-frequency coupling (CFC) spectrograms (Fig. 1B) to confirm electrodes targeted the CA1 stratum pyramidale [43]. Local field potentials (LFPs) were recorded either in a silent background or in the presence of loud broadband noise during locomotion on the treadmill (maximal speed 12 cm/s) (Fig. 1C). The Shapiro-Wilk test confirmed a normal distribution of theta frequency (7-10 Hz) during both silence (w=0.969, p=0.677) and noise (w=0.959, p=0.452). However, the power distribution of theta oscillations did not fit a normal distribution (silence: w=0.891, p=0.016; noise: w=0.788, p=0.000). Analysis of the mean power spectral density (PSD) of the LFPs showed that loud noise significantly increased the mean theta frequency (silence: 7.68 ± 0.11 Hz; noise: 8.36 ± 0.10 Hz, Wilcoxon test, p<0.0001, n=23, Fig. 1F-G) and decreased theta power (silence: 3650 ± 417.4 μV²/Hz; noise: 2262 ± 309.8 μV²/Hz, Wilcoxon test, p<0.0001, n=23, Fig. 1H). Further analysis of higher frequencies in the LFPs revealed a significant decrease in the mean power of slow gamma oscillations in the noise condition (40-70 Hz; 234.6 ± 48.06 μV²/Hz for the silence protocol, and 198.5 ± 41.77 μV²/Hz for the noise protocol, Wilcoxon test, p=0.0068, Fig. 1H).

**Figure 1.**
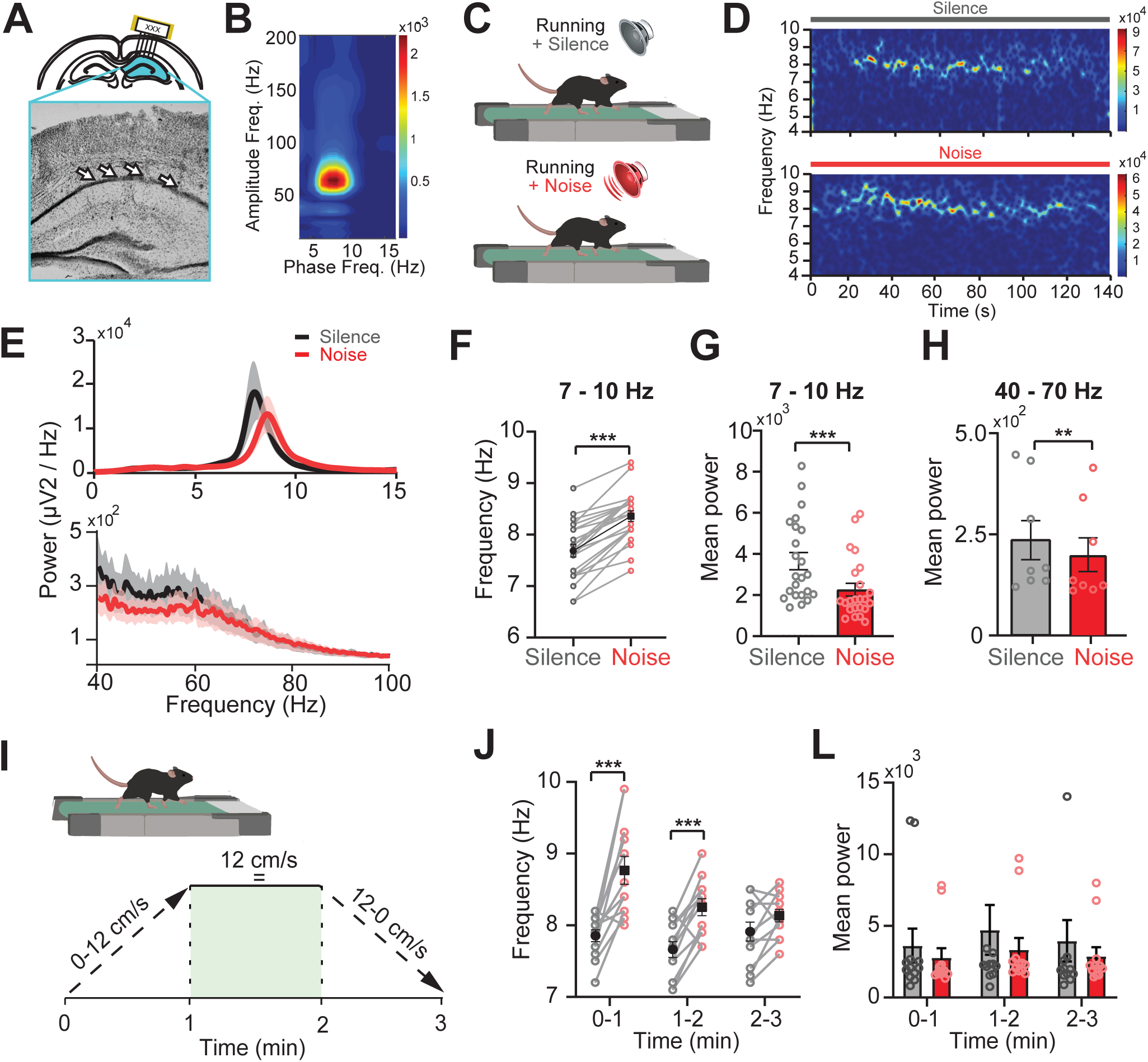
Broadband noise increases theta frequency and decreases theta power during running. A) Position of electrode array into the dHPC (top) and bright-field image of the CA1 with electrode tracts terminating in the stratum pyramidale (arrows). B) Cross-frequency coupling spectrogram of the LFP confirming electrode location in the CA1 *stratum pyramidale*. C) The treadmill task in silence or during noise. D) Representative spectrogram in relation to treadmill speed during silence (top) and during noise stimuli (bottom). E) Mean power spectral density (PSD) plot of the LFP signal (0-15 Hz) (top) during running in silence (gray) and during loud noise (red). Mean PSD of the LFP signal (40-100 Hz) (bottom) during running in silence (gray) and during loud noise (red). Lines show mean power density, and shading shows SEM. F) Line graph showing increased locomotion-induced theta of mice during silence (gray) or noise stimuli (red) (p=0.0001, n=23 mice). H) Bar graph showing a significant decrease in mean theta power during noise compared to silence (p=0.0001, n=23 mice). J) Bar graph showing decreased mean slow gamma power compared to silence (p=0.0068, n=7 mice, Wilcoxon test, **p≤0.001). I) Illustration of the linear changes in speed vs. time on the treadmill. J) Firing frequency at three different intervals (0-1 min, 1-2 min, and 2-3 min) during treadmill running. Black squares show mean values, silence (gray circles), and noise (red circles). During the first and second minutes, firing frequency increased in response to noise (p=0.0005 and p=0.0004, respectively, n=11). L) Bar graph showing no change in mean LPF power for the three running intervals, silence (gray), noise (red) , n=11. One-way repeated measures ANOVA -Tukey’s HSD test, error bar shows SEM, ***p≤0.001. Error bars represent the standard error of the mean (SEM).

We repeated the experiment with another set of animals (n=11) using a treadmill with a variable speed protocol to assess whether the changes in oscillation frequency and power depended on speed and acceleration on the treadmill. In this protocol, the speed increased linearly from 0 to 12 cm/s over the first minute, remained constant at 12 cm/s for the second minute, and decreased linearly from 12 to 0 cm/s during the third minute. The distribution was normal for all of theta frequency (7-10 Hz) protocols, during the first minute, there was a significant increase in frequency during noise stimuli compared to silence (silence: 7.85 ± 0.08 Hz; noise: 8.76 ± 0.19 Hz, T-test, p=0.0005, n=11, Fig. 1J), and between 1-2 min, a significant increase in frequency under noise compared to silence (silence: 7.65 ± 0.11 Hz; noise: 8.25 ± 0.11 Hz, T-test, p=0.0004, n=11, Fig. 1L). However, there was no significant difference in frequency between noise and silence conditions at the last minute. No significant differences were observed in the mean power between silence and noise conditions across the time intervals (0-1 min: p=0.1099, 1-2 min: p=0.3394, 2-3 min: p>0.9999, Wilcoxon test, n=11).

To assess whether any improvement in animal performance in the treadmill could interfere with the noise response, we repeated the experiment with other animals (n=10) for four consecutive days and varied the order of stimuli presentation (Supplementary Fig. 1). We found results to be robust as animals showed an increase in the type 1 theta frequency during noise stimuli at each day: day 1 (silence: 8.250 ± 0.1996 Hz; noise: 8.700 ± 0.1291 Hz, T-test, p=0.001); day 2 (silence: 8.100 ± 0.1291 Hz; noise: 8.533 ± 0.1145 Hz, T-test, p=0.035); day 3 (silence: 8.017 ± 0.2428 Hz; noise: 8.583 ± 0.1701 Hz, T-test, p=0.003); day 4 (silence: 8.183 ± 0.1138 Hz; noise: 8.550 ± 0.0847 Hz, T-test, p=0.003, Supplementary Fig.1B). We also tested if pure tone stimulation could modify theta oscillations but found no significant difference between the peak frequency of theta between silence (8.200 ± 0.1980 Hz) and a 20 kHz pure tone stimulus (8.271 ± 0.39 Hz, p=0.1629), (Wilcoxon Test, p=0.3750, n=8, Supplementary Fig.1 C-D). Together our results show that dHPC locomotion-induced oscillations are specifically sensitive to loud broadband noise.

### Noise modulation of dHPC theta frequency depends on entorhinal input and is not modulated by auditory cortex activity

To investigate if the dHPC response to broadband noise is modulated by noise processing by the auditory cortex, we inhibited pyramidal neurons of the primary auditory cortex (A1) using chemogenetic viral vectors expressing mutated muscarinic M4 receptors (hM4Di) linked to the Ca^2+^/calmodulin kinase 2 alpha (CaMKIIL) promoter (pAAV9-CaMKIIL-hM4D(Gi)-mCherry). Mice (n=4) were bilaterally injected with CaMKIIL-hM4Di-mCherry viral vectors at the time of electrode implants and recordings were carried out 3-4 weeks later to ensure adequate expression of hM4Di receptors, also confirmed post hoc by visualization of mCherry at the injection site (Fig. 2A). On the experimental day mice were injected i.p. with saline (control) or CNO (0.5 mg/kg, chemogenetic inhibition) 10 or 40 min, respectively, prior to the treadmill task during auditory stimulation. The LFP was recorded in the dHPC while animals were running on the treadmill in four different sessions (Saline+silence, Saline+noise, CNO+silence, and CNO+noise; Fig. 2B). The Mauchly’s test was used for validation of repeated measures analysis of variance (ANOVA) for theta frequency data, and the sphericity was assumed χ^2^(5)=8.923, p=0.133 and the distribution was significantly normal in all recordings (Saline+silence; w=0.8715, p=0.303; Saline+noise: w=0.919, p=0.492; CNO+silence: w=0.949, p=0.712, CNO+noise: w=0.828, p=0.162). We found a significant difference in the theta frequency [F(3, 3) =31.92, p=0.0001] when comparing the different recording sessions using One-way repeated measures ANOVA (Fig. 2C). More specifically, Tukey’s HSD test for multiple comparisons found that the theta frequency score was significantly different between silence (6.875 ± 0.23 Hz) compared to noise (8.35 ± 0.3 Hz, p=0.0003, 95% confidence interval (C.I.)=[-2.135, -0.8146] and CNO+noise (8.2 ± 0.26 Hz, p=0.0005, 95% C.I.=[-2.060, -0.7396]) or CNO+silence (6.82 ± 0.27 Hz) compared with noise (p=0.0002, 95% C.I.=[-0.8646, -2.185] and CNO+noise (p=0.0004, 95% C.I.=[-2.110, -0.7896]; Fig. 2C). There was no statistically significant difference in theta frequency between noise stimuli and CNO+noise stimuli, p=0.983, or silence and CNO+noise, p=0.995, (Fig. 2C). This indicates that the primary auditory cortex does not contribute to the loud noise modulation of theta frequency.

**Figure 2.**
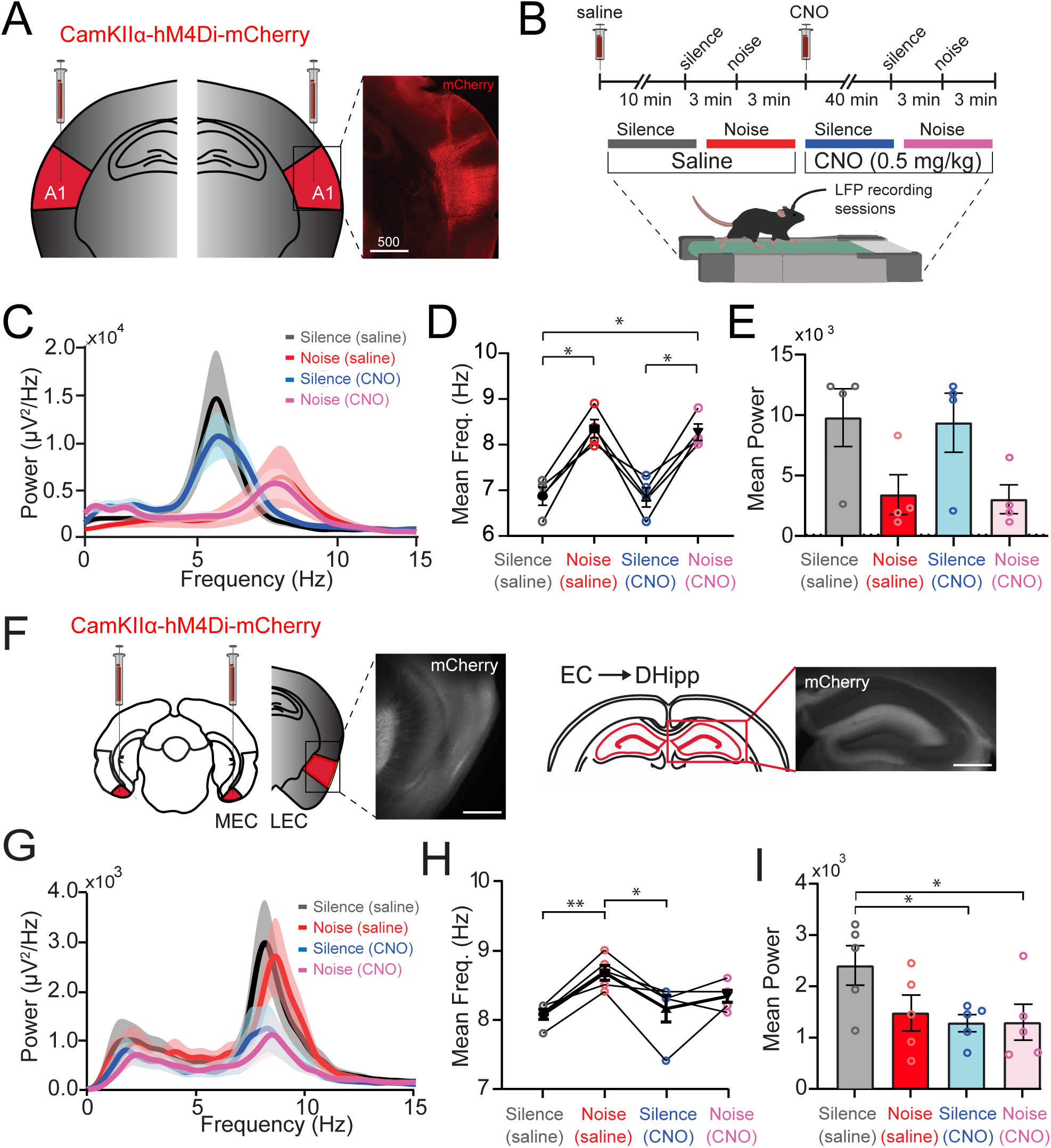
Broadband noise increases theta frequency during locomotion independently of primary auditory cortex activity but requires EC-dHPC pathway activity: A) Illustration of bilateral A1 injections of inhibitory chemogenetic constructs (pAAV9-CaMKIIL-hM4D(Gi)-mCherry). *Inset:* Fluorescence image showing mCherry expressing pyramidal neurons of the A1 region. B) Experimental design of silence and sound sessions after administration of saline and CNO. C) PSD plot of LFP signal (0-15 Hz) recorded 10 min after a saline i.p. injection during silence (gray), noise stimuli (red), and 40 min after a CNO injection (0.5mg/kg i.p.) during silence (blue) and noise (pink). D) Mean frequency of theta oscillations significantly differed between silence and noise stimuli, both after saline and CNO injections (p=0.01, n=4 mice). E) Mean theta oscillation power between silence and noise independently of saline or CNO injections. n=4 mice, Friedman test, Error bars show SEM, *p < 0.05, ***p < 0.001. F) Schematic of bilateral injections of pAAV9-CaMKIIL-hM4D(Gi)-mCherry into the medial and lateral entorhinal cortex (red regions; MEC, LEC). The fluorescent image shows mCherry (white, for better contrast) in the LEC. *Bottom:* Projections for the EC terminating in the dHPC visualized by mCherry (white, for better contrast). G) Mean PSD plot of the LFP signal (0-15Hz) recorded as in ‘C’ H) Line graph depicting the mean theta 1 frequency response under the same conditions as in panel ‘G’. Significant differences in mean theta 1 frequency were observed between silence (saline) and noise (saline) and between silence+CNO (pink) and noise+CNO (blue). I) Bar graphs showing the difference in mean power density between saline+silence and CNO+silence (blue), and after CNO+noise (pink). One-way repeated measures ANOVA -Tukey’s HSD test, error bar shows SEM, *p<0.05, **p<0.01.

For power spectrum analysis, since sphericity was violated (χ^2^(5) =19.206, p=0.005, ε=0.339 using Mauchly’s test), we used Geisser-Greenhouse correction. The distribution was not normal for all protocols (silence: w=0.712, p=0.016; noise: w=0.785, p=0.078; CNO+silence: w=0.678, p=0.006, CNO+noise: w=0.837, p=0.188). Since the data was not normally distribute,d a non-parametric Friedman test of multiple comparisons was used and revealed a significant (p=0.015) decrease in the theta power during the noise stimuli (3096 ± 2092 Hz μV^2^/Hz) compared to silence (9522 ± 2788 μV^2^/Hz; Fig. 2D). However, no statistically significant difference in theta power scores were seen between Saline+silence and CNO+silence (8876 ± 3394 μV^2^/Hz, p=0.999), or Saline+silence and CNO+noise (2958 ± 1743 μV^2^/Hz, p=0.082), or Saline+noise and CNO+silence (p=0.3314), or Saline+noise and CNO+noise (p=0.999), or CNO+silence and CNO+noise (p = 0.999, Fig. 2D). This indicates that inhibiting/decreasing firing frequency in the primary auditory cortex in response to broadband noise does not significantly alter the power of the dHPC theta oscillation during locomotion. Together, bilateral chemogenetic inhibition of the A1 does not interfere with loud broadband noise increase of locomotion-induced theta 1 (7-10 Hz) frequency of the dHPC, thereby suggesting the effect to be due to a non-canonical pathway for sound processing.

To investigate the contribution of the entorhinal cortex inputs to the dHPC we injected both the medial entorhinal cortex (mEC) and the lateral entorhinal cortex (LEC) of C57BL/6 mice bilaterally (n=5) with inhibitory chemogenetic constructs (AAV9.CaMKIIa.hM4D(Gi).mCherry) during the electrode implant surgery and confirmed hM4Di-mCherry expression post hoc (Fig. 2F). The LFP was recorded in the dHPC while the animals were running on the treadmill in four different sessions, 10 min after an i.p. injection with saline (control, silence and noise) and 40 min after the i.p. injection of CNO (0.5 mg/kg, silence and noise). For theta frequency data, the sphericity was assumed χ^2^(5) = 2.793, p=0.747 and the distribution was significantly normal in all recordings (Saline+silence: w=0.778, p=0.053; Saline+noise: w=0.973, p=0.898; CNO+silence: w=0.648, p=0.055; CNO+noise w=0.953, p=0.758). We found a significant difference in the theta frequency [F(3, 4)=7.863, p=0.004] when comparing the different recordings sessions, where Tukey’s HSD test for multiple comparisons shows the mean theta frequency score to be significantly different between Saline+silence (8.01 ± 0.11 Hz) and Saline+noise (8.68 ± 0.18 Hz, p=0.004, 95% C.I.=[-0.9991, -0.2009]), and for Saline+noise and CNO+silence (p=0.010, 95% C.I.=[-0.1209, -0.9191], Fig. 2G-H). However, there was no difference in theta frequency in runs comparing Saline+silence (8.01 ± 0.11 Hz) and CNO+silence (8.16 ± 0.30 Hz, p=0.932) or Saline+silence and CNO+noise (8.3 ± 0.15 Hz, p=0.265), or noise and CNO+noise (p=0.1052), or CNO+silence and CNO+noise (p=0.5514, Fig. 2G-H). Thereby it appears that i.p CNO administration, to inhibit excitatory input from the medial and lateral entorhinal cortex, removed the broadband noise effect of increasing theta 1 frequency. Thus, the entorhinal cortex appears to contribute to loud broadband noise modulation of dHPC locomotion-induced theta 1 oscillations.

For power spectrum analysis, the sphericity was assumed χ^2^(5) =5.937, p=0.339 and the distribution was significantly normality to all recordings (Saline+silence: w=0.904, p=0.434; Saline+noise: w=0.967, p=0.857; CNO+silence: w=0.886, p=0.340, CNO+noise: w=0.851, p=0.198). We found a significant difference in the theta power score [F(3, 4)= 4.801, p=0.020] when comparing the animals in the different sessions. More specifically, Tukey’s HSD test for multiple comparisons found that the theta power score was significantly different between silence (2411 ± 699 μV^2^/Hz) compared with CNO+silence (1280 ± 300 μV^2^/Hz, p=0.029, 95% C.I.=[-102.9, -2160]) and silence compared to CNO+noise (1299 ± 565 μV^2^/Hz, p=0.032, 95% C.I.=[-83.30, -2141]; Fig. 2I). However, there was no statistically significant difference in theta power scores between silence and noise (1479 ± 612 μV^2^/Hz, p=0.080) or Saline+noise and CNO+silence (p=0.937), or Saline+noise and CNO+noise (p=0.953), or CNO+silence and CNO+noise (p=0.999, Fig. 2I). Together these results indicate that the entorhinal cortex is important for mediating loud noise in the environment contributing to the increase in theta 1 frequency during running, while the entorhinal cortex inputs contribute to theta oscillation power magnitude during silence but not during loud noise.

### Optogenetic stimulation of DCN neurons that project to PRN increase theta frequency and decrease theta power in the dHPC

Sound is first processed in the cochlear nucleus, independently if going through the canonical or non-canonical pathway of sound processing [8]. The dorsal cochlear nucleus projects collaterals via de pontine reticular nucleus (PRN), a nucleus important for startle responses [44,45]. To investigate if the DCN-PRN pathway also contributes to loud noise increase in mean theta frequency, C57BL/6 mice (n=3) were bilaterally injected with retrograde viral particles (AAVrg-CAG-hChR2-H134R-tdTomato) into the PRN (Fig. 3A), and bilaterally implanted with optic fiber ferrules, with fiber tips reaching the dorsal border of the DCN (Fig. 3B), as well as the electrode array implanted in the dHPC. On the day of the experiment, animals were connected to the headstage and dual optic fibers were connected to one blue laser source (488 nm), and the animal was placed on the treadmill. The LFP was recorded in the dHPC while the animals were running on the treadmill in three sessions: silence, noise, and blue light “noise” (frequency and amplitude modulated waveform of randomized components, see methods) to activate channelrhodopsin expressing DCN neurons projecting through the pontine reticular nuclei. Optogenetic stimulation during locomotion elicited change in both the mean theta frequency and power of theta oscillations in a manner similar to loud broadband noise modulation of theta (Fig. 3B). For theta frequency data, the sphericity was assumed χ^2^(2)= 3.348, p=0.187 (Mauchly’s test) and the distribution was normal for all recordings (silence: w=0.986, p=0.7804; noise: w=0.9796, p=0.7262; blue light: w=0.7742, p=0.0542). We found a significant difference in the theta frequency [F(2, 2) =23.79, p=0.006] when comparing all recordings using One-way repeated measures ANOVA. More specifically, Tukey’s HSD Test for multiple comparisons found that the theta frequency score was significantly different between silence (7.833 ± 0.1453 Hz) and noise (8.633 ± 0.2333 Hz, p=0.0052, 95% C.I.=[-1.215 to -0.3849]) and blue light stimuli (8.297 ± 0.1017 Hz, p=0.0353, 95% C.I.=[-0.8784 to -0.04827], Fig. 3C-D). Interestingly, there was no statistically significant difference in locomotion-induced theta frequency in sessions of noise and blue light “noise” of the DCN-RPN pathway (p=0.100, Fig. 3D). This suggests that broadband noise is processed in the non-canonical pathway and that optogenetic stimulation of DCN neurons projecting through the RPN can modulate dHPC theta frequency, but not as strongly as loud broadband noise stimuli.

**Figure 3.**
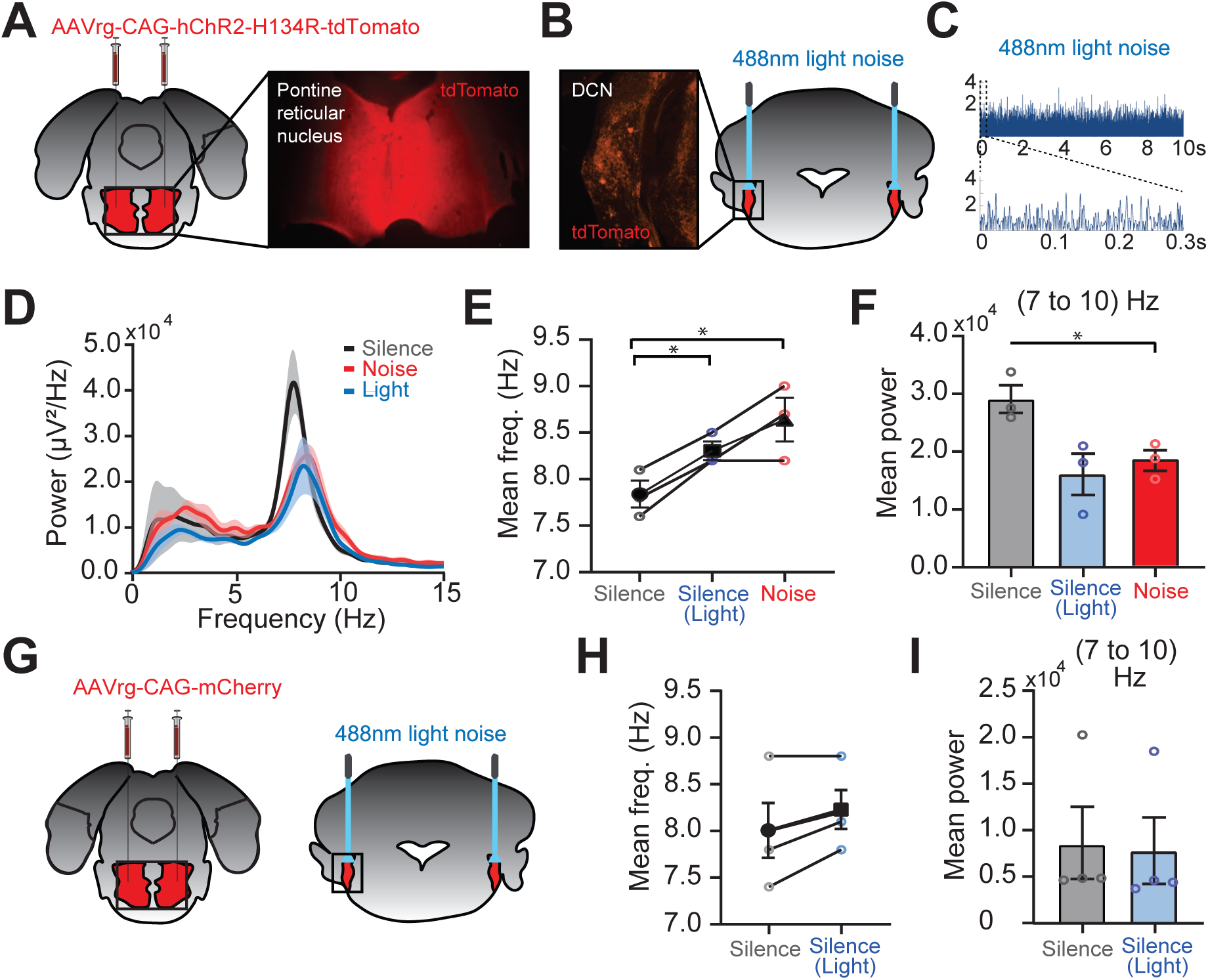
Optogenetic stimulation of DCN neurons projecting to the PRN increases theta frequency and decreases theta power in the dHPC. A) Schematic of bilateral injections into the pontine reticular nucleus (PRN, red) with a retrograde optogenetic construct (AAVrg-CAG-hChR2-H134R-tdTomato). Fluorescent image showing tdTomato indicating channelrhodopsin 2 (ChR2) expression in axonal terminals in the PRN. B) *Left*: Fluorescence image showing tdTomato expressing DCN neurons, indicating that they project to the PRN. *Right:* Schematic drawing of bilateral optic fiber implants into the most dorsal part of the DCN. C) Light (473 nm) stimuli mimicking a broadband noise (see methods). D) Mean PSD plot of LFP signal (0-15Hz) recorded as mice (n=3) run on a treadmill during silence (gray), blue light stimulation (blue), and noise (red). E) Line graph showing individual theta 1 frequency response of the dHPC during silence (gray), optogenetic stimulation (blue), and noise stimuli (red). The mean theta 1 frequency was significantly different between silence (circle) and optogenetic stimulation (square) and between silence and noise stimuli (triangle), p=0.0392 and p=0.0303, respectively (n=3). F) Bar graph showing decreased mean theta power during noise stimuli (red) and optogenetic stimulation (blue) compared to during silence (gray), p=0.0052 and p=0.0353, respectively, (n=3). One-way repeated measures ANOVA, error bars show SEM, *p<0.05, **p<0.01. G) Schematic of control experiment with retrograde constructs (AAVrg-CAG-mCherry) injected into the PRN and bilateral optic fiber implants into the most dorsal part of the DCN. H) Line graph showing no change in theta due to light stimulation silence (gray), and light noise (blue). F) Bar graph showing no difference in mean theta power during silence (gray) and light stimulation (blue) in mice not expressing ChR2. T-test, error bars show SEM, *p<0.05, **p<0.01.

For power spectrum analysis of optogenetic stimulation of the DCN-PRN pathway, the sphericity was assumed χ^2^(2) = 0.912, p = 0.634, and the distribution was significantly normality to all recording sessions (silence: w=0.8987, p=0.3813; noise: w=0.9913, p=0.8213; blue light stimulation w=0.9188, p=0.4482). First, a significant difference in the theta power [F (2, 2)=23.79, p=0.0060] was found when comparing all recordings. More specifically, Tukey’s HSD test for multiple comparisons showed that the theta power density was significantly different between mice running on the treadmill in silence (2907 ± 239.5 μV^2^/Hz) compared to during loud noise stimuli (1847 ± 177 μV^2^/Hz, p=0.0052, 95% C.I. = [-1.215 to -0.3849], Fig. 3E), and compared to during optogenetic stimulation (1608 ± 355.9 μV^2^/Hz), p=0.0353, 95% C.I. = [-0.8784 to -0.04827], Fig. 3E). There was no statistically significant difference in theta power scores between broadband noise and blue light stimuli (p=0.0926, Fig. 3E). Together this indicates that the activation of the non-canonical DCN-PRN pathway can increase mean theta frequency, and decrease theta power, during locomotion similarly to when exposing mice to loud broadband noise during running.

### Broadband noise increases theta frequency and activates MS PV+ interneurons independently of MS cholinergic activity

To investigate the influence of MS cholinergic neurons on the theta response to broadband noise, we decreased activity of cholinergic transferase (ChAT) expressing neurons in the MS by injecting ChAT-cre transgenic mice (n=6) with inhibitory chemogenetic viral vectors dependent on cre recombinase activity (pAAV9.hSyn.DIO.hM4D(Gi).mCherry) into the medial septum (Fig. 4A) during the same surgery as for the dHPC electrode array. Similar experimental sessions as for testing contributions of the A1 and EC were done, where the dHPC LFP was recorded when animals were running on the treadmill in silence and in the presence of broadband noise, after saline and CNO i.p. injections, respectively (Fig. 4B). For theta frequency data, the sphericity was assumed χ^2^(5) =2.793, p=0.747 and the distribution was significantly normal to all recordings (Saline+silence: w=0.975, p=0.924; Saline+noise: w=0.759, p=0.052; CNO+silence: w=0.935, p=0.625, CNO+noise: w=0.845, p=0.143). A significant difference in the theta frequency [F(3, 5)=8.527, p=0.0015] was found when comparing different recording sessions (Fig. 4C). Specifically, Tukey’s HSD test for multiple comparisons showed the mean theta frequency score was significantly different between silence (7.26 ± 0.2044 Hz) compared with noise (7.837 ± 0.1554 Hz, p=0.0191, 95% C.I.=[-1.056 to -0.08431]), or silence and CNO+noise (7.883 ± 0.1956 Hz, p=0.011, 95% C.I.=[-1.102 to -0.1310]) or CNO+silence (7.250 ± 0.2655 Hz) and noise (p=0.0158, 95% C.I.=[0.1010 to 1.072]) or CNO+silence and CNO+noise (p=0.0092, 95% C.I.=[-1.119 to - 0.1476], Fig. 4C). Thereby CNO application had no significant effect on locomotion-induced theta frequency as there was no difference between the session in Saline+silence and CNO+silence (p=0.999) or sessions with noise and CNO+noise (p=0.990, Fig. 4C).

**Figure 4.**
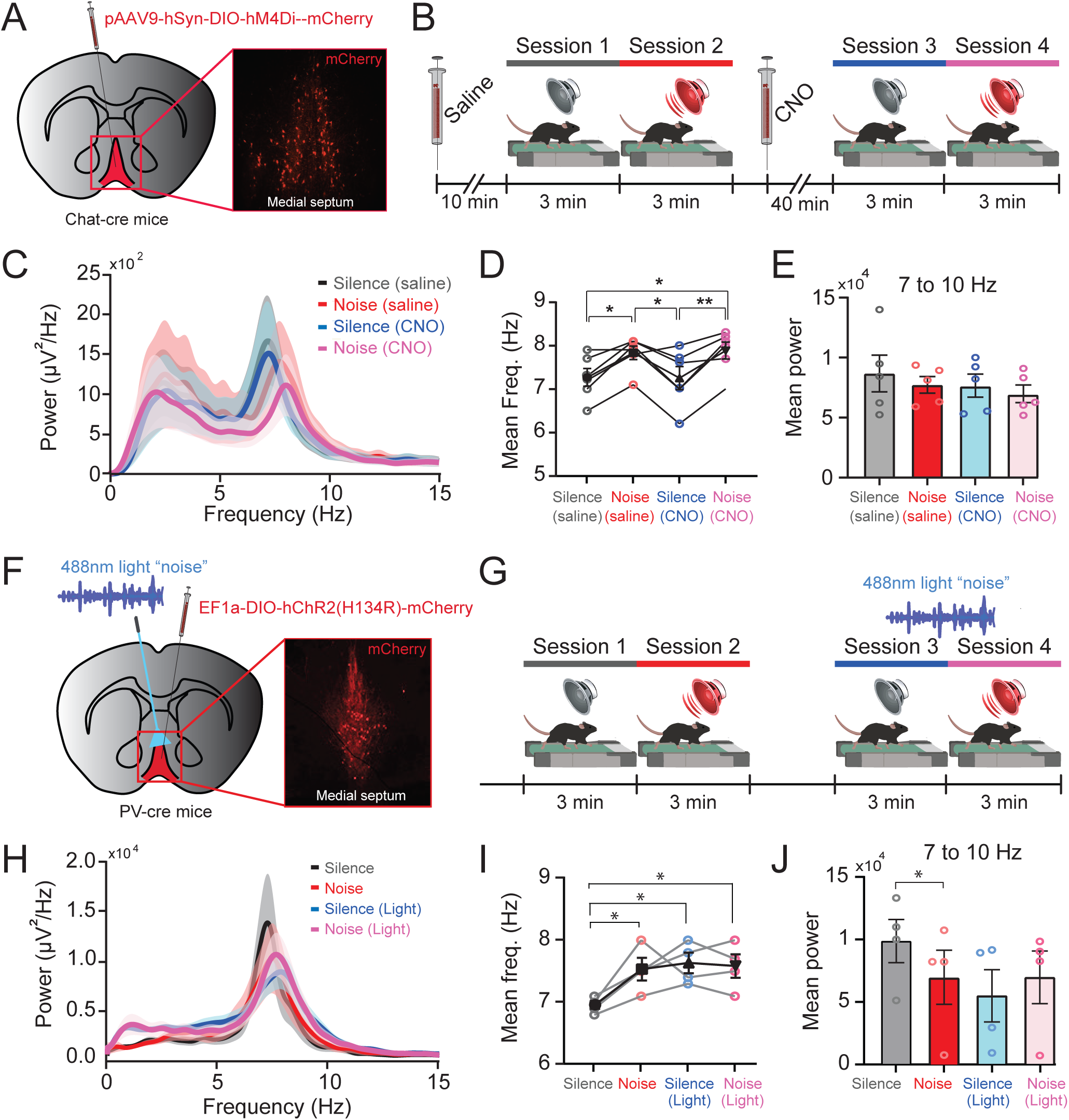
Noise stimuli require MS PV+ neuron activity but not cholinergic activity for increasing dHPC theta frequency. A) Schematic drawing of the angled (12 degrees) injection of cre-dependent chemogenetic construct (pAAV9.hSyn.DIO.hM4D(Gi).mCherry) into the MS of Chat-cre mice (n=5). *Right,* Fluorescent image of mCherry expression in Chat+ neurons of the MS. B) Experimental schematic of silence and sound sessions after administration of saline and CNO. C) Mean PSD plot of the LFP (0-15Hz) of the dHPC during treadmill running upon saline and CNO injection (0.5mg/kg, i.p.) during silence and noise. D) Line graph showing mean theta 1 frequency is significantly different between silence (gray and blue) and noise stimuli (red and pink) independent of saline or CNO injection *p<0.05, **p<0.01. E) Bar graph shows that the mean power was not significantly different when inhibiting MS Chat+ neurons independently of stimuli or treatment. F) *Left;* Schematic drawing of optic fiber inserted with a 12-degree angle into the MS of PV-cre mice (n=4) previously injected with a cre-dependent optogenetic construct for channelrhodopsin expression (pAAV9-EF1a-double floxed-hChR2(H134R)-mCherry). *Right:* Fluorescent image showing mCherry expression by PV+ MS neurons. G) Mean PSD plot of the dHPC LFP signal (0-15Hz) recorded during silence (gray), noise (red), light (473 nm, light noise, see methods, blue), and blue light during noise stimuli (pink). H) Mean frequency of theta frequency was significantly different between silence and noise (red), light stimulation (blue), and light+noise stimuli (pink). I) Mean power was higher for silent recording than noise (p=0.0411 Friedman test of multiple comparisons). Error bars show SEM, *p<0.05.

For power spectrum analysis, the sphericity was assumed χ^2^(5) = 8.923, p=0.133 and the distribution was significantly normality to all recordings (Saline+silence: w=0.931, p=0.606; Saline+noise: w=0.894, p=0.382; CNO+silence: w=0.879, p=0.304; CNO+noise: w=0.911, p=0.479). There was no statistically significant difference in theta power scores [F(3, 4)=2.171, p=0.1445] between silence and noise after saline (p=0.5127), or Saline+silence and CNO+silence (p=0.4561), or silence and CNO+noise (p=0.1044), or noise and CNO+silence (p=0.999), or Saline+noise and CNO+noise (p=0.686), or CNO+silence and CNO+noise (p=0.745, Fig. 4D). Thereby, cholinergic MS neurons do not appear to modulate loud noise induced increase in mean theta frequency or decrease in theta oscillation power.

To investigate the influence of Parvalbumin-positive MS neurons on the dHPC response to broadband noise, we wanted to control the activity of inhibitory PV neurons using channelrhodopsin2 during sessions of locomotion in silence or in the presence of broadband noise. Thereby, PV-cre positive transgenic mice (n=4) were injected with viral vectors (pAAV9-EF1a-double floxed-hChR2(H134R)-mCherry) into the MS together with the implant of a fiber optic ferrule with the fiber tip reaching the MS in a 12° angle (Fig. 4F) as well as implanting the electrode array. The dHPC LFP was recorded when animals were running on the treadmill in silence and in the presence of broadband noise, with or without concomitant blue light stimulation (Fig. 4G). For theta frequency data, the sphericity was assumed χ^2^(5)=2.527, p=0.805, and the distribution was significantly normal to all data (silence: w=0.863, p=0.272; noise: w=0.938, p=0.647; silence+light w=0.915, p=0.513, noise+light w=0.993, p=0.976). We found a significant difference in the theta frequency [F(3, 3)=6.378, p=0.0132] when comparing all recordings (silence: 6.950 ± 0.06455 Hz, noise: 7.5225 ± 0.1843 Hz, silence+light 7.625 ± 0.1652 Hz, and noise+light 7.575 ± 0.1887 Hz) using One-way repeated measures ANOVA (Fig. 4H). More specifically, Tukey’s HSD Test for multiple comparisons found the mean theta frequency score significantly different between silence compared with noise stimuli (p=0.0407, 95% C.I.=[-1.126 to -0.02406]), silence+light stimulation, p=0.0175, 95% C.I.=[-1.226 to -0.1241] and noise+light, p=0.0267, 95% C.I.=[-1.176 to -0.07406], Fig. 4H). However, there was no statistically significant difference in mean theta frequency between noise stimuli and silence+light (p=0.9395 or noise and noise+light (p=0.9915) or silence+light and noise+light (p=0.9915, Fig. 4I). This indicates that, during running, inhibition provided by PV+ MS neurons has a similar effect as loud noise on dHPC theta oscillations. Interestingly, PV+ MS neuron activation did not further increase the mean theta frequency upon loud noise stimulation (Fig. 4I).

For power spectrum analysis, since sphericity was violated χ^2^(5) =13.268, p=0.041, ε=0.386, we used Geisser-Greenhouse correction. The distribution was not significantly normal for all recordings (silence: w=0.865, p=0.1357; noise: w=0.756, p=0.009; silence+light: w=0.765, p=0.012, noise+light: w=0.7016, p=0.002) yet using a Friedman test of multiple comparisons could reveal a significant difference (p=0.022) in the theta power between silence (9940 ± 1737 μV^2^/Hz) and noise (7036 ± 2174 μV^2^/Hz, Fig. 4J). However, there was no statistically significant difference in theta power between silence and silence+light (5540 ± 2108 μV^2^/Hz, p=0.071) or silence and noise+light (7022 ± 2122 μV^2^/Hz, p=0.120), or noise stimuli and silence+light (p=0.999) or noise and noise+light stimulation (p=0.999), or silence+light stimulation and noise+light stimuli (p=0.999, Fig. 4J). This indicates that dHPC theta power is not modulated by PV+ interneurons of the MS, however, recordings showed large variability in mean theta (7-10Hz) power in these animals. Taken together, PV+ interneurons of the MS appear to modulate theta frequency, but not theta power, during running independently if mice are running in a silent environment or in the presence of loud noise.

### dHPC pyramidal neuron calcium activity is reduced during treadmill running in loud noise compared to in silence

As broadband noise consistently showed an increase in theta frequency but a reduction in power, we wanted to measure the calcium activity of CA1 pyramidal neurons in the dHPC in response to loud noise and if activating the DCN neurons of the DCN-PRN pathway could mimic noise response in the dHPC. Therefore, mice (n=3) were injected with retrograde expressing channelrhodopsin2 in the PRN, to express ChR2 in DCN neurons connected to the PRN, and also with CaMKII-GCaMP6f into the dHPC CA1 region (Fig. 5A). Channelrhodopsin2 expressing neurons of the DCN were excited bilaterally and since the DCN is a more caudal structure than the dHPC, the optic fiber ferrules did not impede the fitting of the miniscope when mice were running on the treadmill (Fig. 5B). Expression of GCaMP6f in CA1 *stratum pyramidale* was confirmed post hoc (Fig. 5B) and calcium dynamics was extracted from 661 CaMKII+ pyramidal neurons from recordings when mice were running on the treadmill in three sessions: silence, during loud broadband noise and silence during blue laser stimulation (Fig. 5C-D).

**Figure 5.**
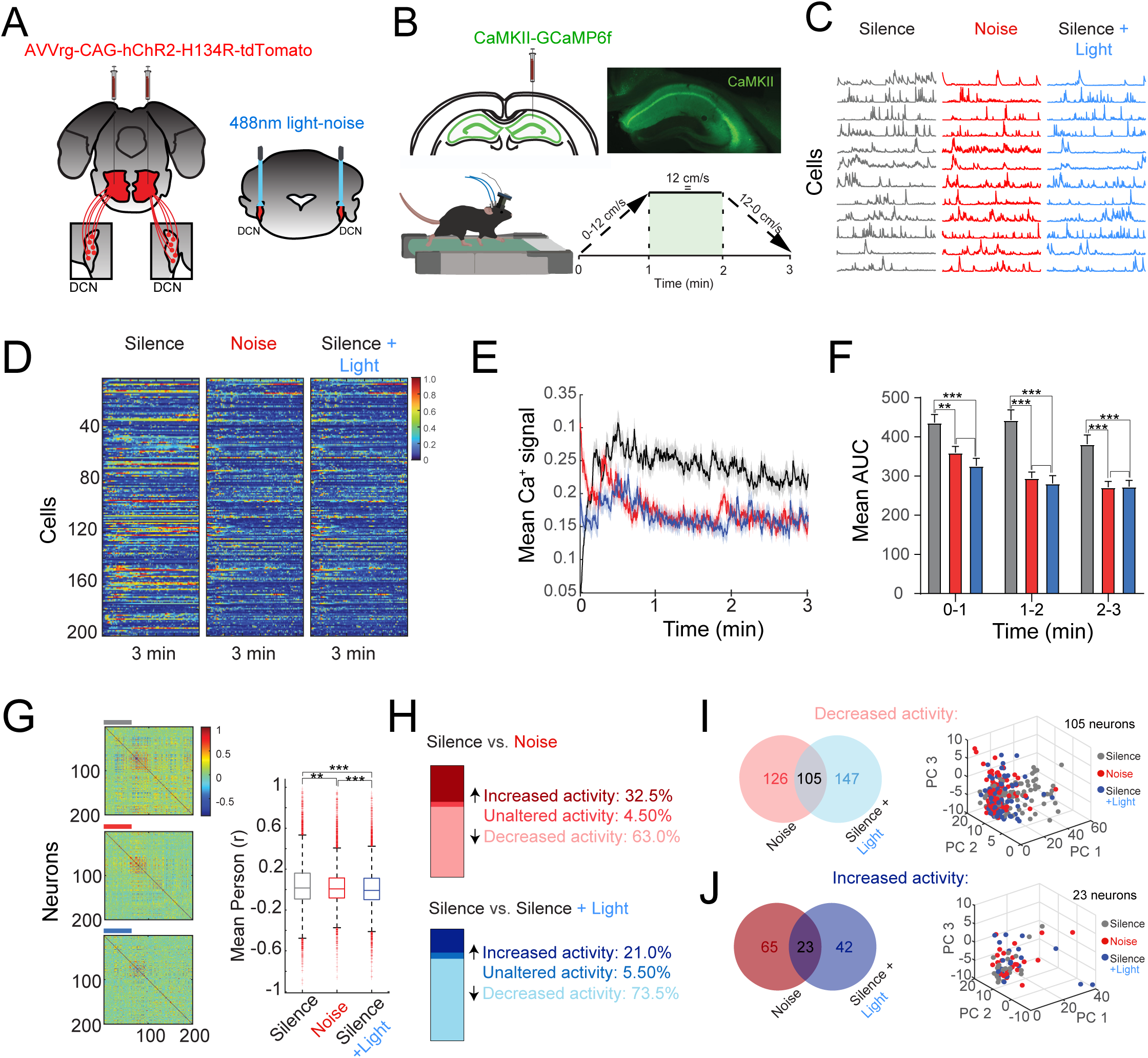
Noise stimulation activates the reticulo-limbic pathway that reduces CA1 pyramidal cell activity of the dHPC. A) Schematic drawing of retrograde channelrhodopsin2 expression in the DCN. B) Experimental design shows CaMKII-GCaMP6f expression in the dHPC (*top*) and miniscope recording during treadmill running C) Representative calcium dynamics traces from individual CaMKII+ neurons during silence (gray), noise (*red*), and light stimulation (blue). D) Heat maps showing normalized calcium activity of 200 neurons over a 3-minute period under three conditions: Silence (gray), noise (red), and light stimulation (blue). Each row represents the activity of a single neuron, and the color intensity corresponds to the activity level. E) Plot of the mean calcium signal over time for CaMKII-GCaMP6f expressing neurons during silence (black), noise (red), and silence combined with light stimulation (blue). The shaded areas represent SEM. F) Bar graphs showing the mean area under the curve (AUC) for the calcium signals in three consecutive 1-minute intervals (0-1 min, 1-2 min, 2-3 min). Error bars represent the SEM. The mean AUC significantly decreases under Noise and Silence + Light conditions compared to Silence across all intervals (***p < 0.001, n=3). G) *Left,* Correlation matrices showing pairwise correlation coefficients of calcium activity between 200 neurons under three conditions: silence (top), noise (middle), and silence combined with light stimulation (bottom). Color scale: correlation values from -1 to 1. *Right*: box plots comparing neuronal activity mean Pearson correlation coefficients (r) across the three conditions. Error bars represent the standard deviation. The Noise condition shows a significantly higher mean Pearson correlation coefficient than Silence and Silence + Light conditions (***p < 0.001). H) Bar charts depicting the percentage of neurons showing increased (dark color), unaltered, or decreased (light color) activity when comparing Silence to Noise (top) and Silence to Silence + Light (bottom). I) *Left:* Venn diagram showing the overlap of neurons with decreased activity under noise (light red) and Silence+Light (light blue) stimuli compared to silence condition. *Right:* 3D scatter plots showing the principal component analysis (PCA) distribution of overlap neurons with decreased activity. J) Venn diagram showing the overlap of neurons with increased activity under noise (dark red) and Silence+Light (dark blue) stimuli compared to silence condition. *Right:* 3D scatter plots of PCA distribution of overlap neurons with increased activity.

We analyzed the mean calcium activity (ΔF/F) of pyramidal neurons (n=200) across three-time points during the task (Fig. 5E-F). In the first minute, the Area Under the Curve (AUC) was significantly reduced during both noise (360.0 ± 15.50) and silence+light (325.9 ± 19.33) conditions compared to silence (436.4 ± 20.74), with p-values of 0.0013 and <0.001, respectively. In the second minute, AUC remained significantly lower during noise (295.4 ± 14.87) and silence+light (281.5 ± 19.34) conditions compared to silence (443.3 ± 25.48), with p<0.001 for both. By the third minute, AUC continued to be significantly reduced during noise (284.4 ± 15.60) and silence+light (289.7 ± 16.20) compared to silence (404.9 ± 24.65), again with p<0.001 for both conditions (Fig. 5F).

Still, noise and blue light stimulation of the DCN-RPN pathway during locomotion showed appeared to decrease pairwise correlation compared to silence, although the correlation value was very low (silence: r=0.0357 versus noise: 0.02876, p=0.0011; silence versus silence+light: r= 0.0125, p<0.0001; and noise versus silence+light, p<0.0001, n=200 cells, Fig. 8G).

The minority of CaMKII+ cells (9/200, 4.50%) did not alter calcium activity during silence or noise, while 63% (126/200 cells) showed a significant decrease in activity during the noise trial compared to silence, while 32.5% (65/200 cells) increased significantly the activity (Fig. 5H, top). Next, we compared the CAMKll cells modulated by blue light stimulation of the DCN-PRN pathway compared to the silent trial and found 5.50% (11/200 cells) did not show a significant difference, while 73.5% (147/200 cells) showed a significant decrease in activity during the noise trial compared to silence, while 21% (42/200 cells) increased significantly the activity (Fig. 5H, bottom). Of the cells that exhibited modulation compared to silence, 191 cells showed changes during noise and 189 during optogenetic stimulation. Of these, 105 cells that decreased activity compared to silence showed the same profile of changes in both conditions (noise and optogenetic stimulation) (Fig 5I, top). In addition, 23 cells that increased activity also maintained this pattern in both conditions (Fig 5I, bottom). Taken together, results show that pyramidal cells of the dHPC CA1 area are modulated by loud broadband noise, probably from the reticulo-limbic pathway.

### Noise increases the interneuron activity of the dHPC during the first minute

To target GABAergic interneurons, mice (n=3) were injected with GCaMP6f with MDLX promoter in the right dHPC stratum pyramidale region, and calcium dynamics were extracted from 56 inhibitory interneurons (Fig. 6A-C). Given the diversity of interneurons in the hippocampus, we employed the K-means clustering algorithm to group neurons with similar activity patterns (Fig. 6D-F). The optimal number of clusters 3, was determined using silhouette analysis, which also served as a measure of clustering quality. Neurons with silhouette values below 0.1 were excluded from the final analysis to ensure robust classification (n=6 neurons) (Fig. 6D). We identified three distinct clusters in the final analysis (n=50 neurons): Cluster 1, containing 26 neurons (52%); Cluster 2, with 6 neurons (12%); and Cluster 3, with 18 neurons (36%) (Fig. 6E-F). Next, we computed the correlation coefficient of all possible cell pairs to observe any synchrony of the circuit during the different conditions. The correlation between neurons in Cluster 1 increased significantly during noise (r = 0.75) compared to silence (r = 0.34), p < 0.0001 (Fig. 6H, left; and Fig. 6I, top). However, there was no significant difference in pairwise correlation between interneurons in silence and during broadband noise in the other clusters (Cluster 2, p =0.1440; Cluster 3, p =0.5795), (Fig. 6I, middle and bottom).

**Figure 6.**
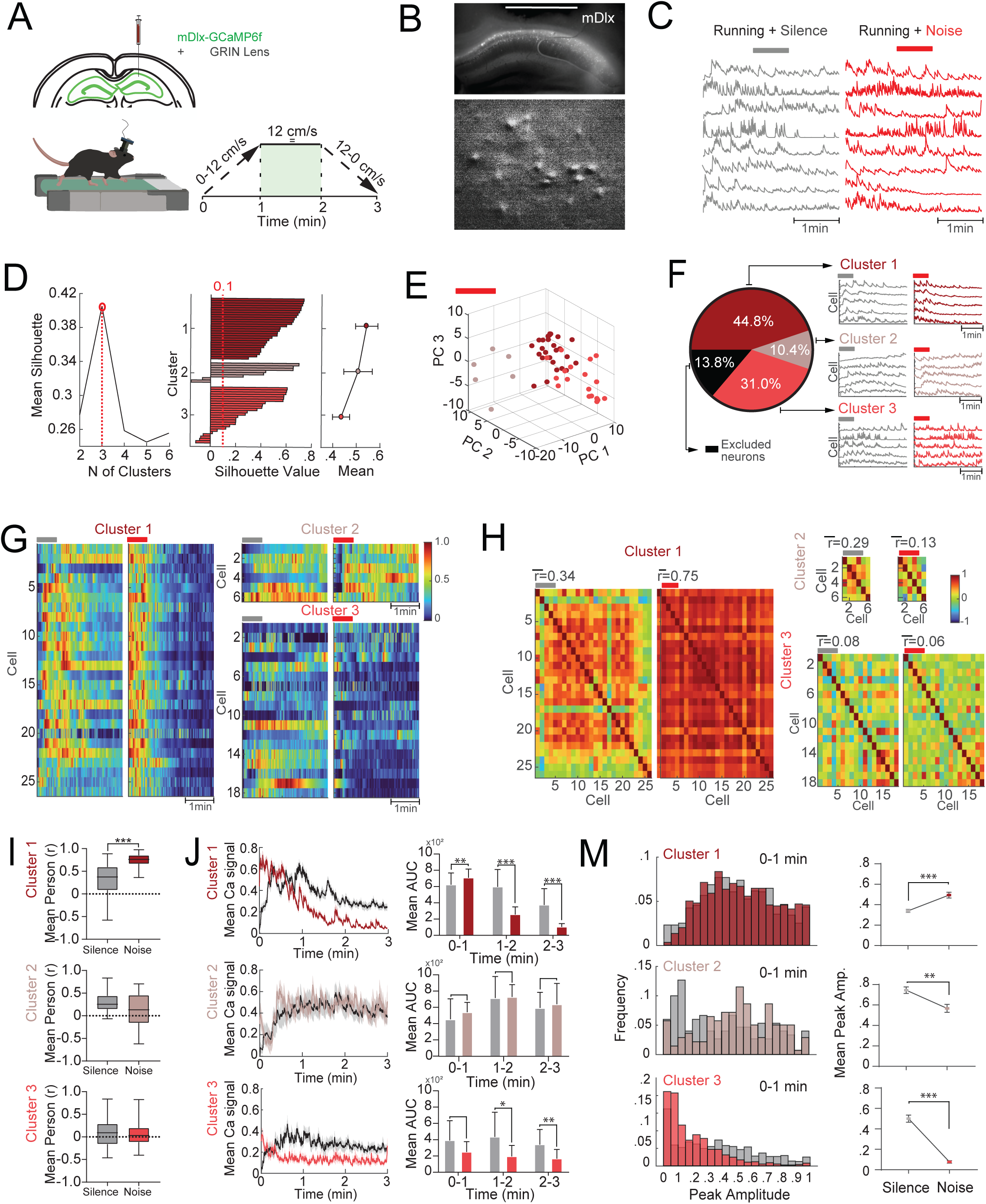
Noise increases interneuron activity of the dHPC. A) Experimental design shows mDlx-GCaMP6f expression in the dHPC (top) and miniscope recording during treadmill locomotion (bottom). B) *Top;* Image of GCaMP6f fluorescence (white) from mDlx positive interneurons of the dHPC CA1 region. *Bottom:* Image from miniscope records shows calcium fluorescent signal (white) from mDlx positive interneurons of the dHPC CA1 region. C) Representative calcium dynamics traces from mDlx+ neurons during silence and noise stimuli. D) Silhouette graph of the optimal number of clusters (Middle) Silhouette plot for 3 cluster solution, red dotted line at 0.1 serves as a threshold for acceptable silhouette values for individual cells. (Right) Mean silhouette values for each cluster with error bars representing standard deviation. E) 3D scatter plot of the distribution of neurons defined by the first three principal components (PC1, PC2, and PC3)from k-means clustering analysis. F) Pie chart of distribution of neurons across the three clusters identified in ‘D’ 13.8% of neurons were excluded from clustering analysis due to their low silhouette scores. (Right) Representative calcium traces for each cluster. Silence (gray), noise (red). G) Heatmap representation of neuronal activity for each cluster. Each row represents a single neuron, with color intensity corresponding to activity level, blue-low activity, red-high activity, during silence (gray bar) and noise (red bar). H) Correlation matrices of neuronal activity within each cluster. The heatmaps represent pairwise correlation coefficients of calcium activity between neurons within each cluster with colors indicating the strength and direction of the correlation (from -1 to 1). I) Box plots of the mean Pearson correlation coefficient distribution for neurons in each cluster under silence (gray) and noise (red) conditions. Only in Cluster 1 the mean correlation significantly increases from silence to noise (***p < 0.001). J) Left: Plots of the mean calcium signal over time for neurons in Cluster 1, 2, and 3. Lines represent mean signals, while shaded areas indicate SEM. Gray - silence and red noise. (Right) Bar graphs showing the mean AUC for the calcium signals in three consecutive 1-minute intervals (0-1 min, 1-2 min, 2-3 min). Error bars represent SEM. In Cluster 1, the mean AUC significantly decreases from the first to the third interval under noise conditions compared to silence (**p < 0.01, ***p < 0.001). In Cluster 3, the mean AUC significantly decreases from the first to the third interval under noise conditions compared to silence (*p < 0.05, **p < 0.01). M) Left: histograms showing the distribution of peak amplitudes for neurons in each cluster during the first minute. Red colored bars - noise, gray bars - silence. Right: bar graphs comparing the mean peak amplitude of calcium signals between silence and noise conditions for each cluster. Cluster 1: The mean peak amplitude significantly increases under noise conditions compared to silence (***p < 0.001), while Cluster 2 and 3 showed a decreased mean peak amplitude compared to silence (**p < 0.01, *p < 0.05, respectively). Error bars show SEM.

Thus, we analyzed the change in the average activity in each cluster in three task times. In Cluster 1, the Area Under the Curve (AUC) increased significantly during the first minute of noise (708.1 ± 20.85), indicating a stronger response compared to silence (622.3 ± 28.47), p = 0.0014. However, in the subsequent minutes two (silence: 596.1 ± 42.11; noise: 256.0 ± 18.31) and three (silence: 373.6 ± 39.47; noise: 103.5 ± 8.019) of the task, activity decreased during noise relative to silence, , p<0.0001 and, p<0.000, respectively (Fig. 6J, top). In Cluster 2, there was no significant difference in mean activity between silence and noise across the three-time points of the task ( min 1: p=0.3702; min 2: p=0.9290, min3: p=0.7896) (Fig. 6J, middle). In contrast, for Cluster 3, the mean activity of the cells decreased during noise compared to silence at two last time points (min 1 - silence: 390.6 ± 57.22; noise: 247.8 ± 30.32, p=0.0563; min 2 - silence: 434.1 ± 71.40; noise: 194.1 ± 31.66, p=0.0189; min 3 - silence: 339.9 ± 43.77; noise: 165.3 ± 27.08, p=0.0049) (Fig. 6J, bottom).

Finally, we analyze the distribution of peak amplitudes for neurons in each cluster during the first minute. Comparing the mean peak amplitude of calcium signals between silence and noise conditions for each cluster. Cluster 1: The mean peak amplitude significantly increases under noise (0.4934 ± 0.02837) conditions compared to silence (0.3396 ± 0.01628) (***p < 0.001) (Fig. 6M, top);, while Cluster 2 (silence: 0.7451 ± 0.03151; noise: 0.5672 ± 0.03825) (Fig. 6M, middle); and cluster 3 (silence: 0.4998 ± 0.03546; noise: 0.07678 ± 0.01159) showed a decreased mean peak amplitude compared to silence (**p <=0.0019, ***p < 0.0001, respectively) (Fig. 6M, bottom).

### Broadband noise modulates mPFC theta oscillation and increases coherence between the dHPC and mPFC

To investigate if broadband noise may influence the modulation of the medial prefrontal cortex (mPFC) theta oscillations [46], we implanted mice (n=10) with custom-made arrays for targeting both the dHPC and the prelimbic mPFC (Fig. 7A). Electrode placement into the mPFC was confirmed with post hoc histology (Fig. 7A) and cross-frequency coupling of the LFP signal confirming the location of the electrode in the mPFC [47] (Fig. 7B). As previously described, mice were placed on the treadmill and the LFP of both the mPFC and the dHPC were recorded during silence and broadband noise stimuli (Fig. 7C). The LFP recordings in the mPFC showed theta frequency distribution to be normally distributed (silence: w=0.941, p=0.571; noise: w=0.915, p=0.316) as was also theta power (silence: w=0.947, p=0.644; noise: w=0.920, p=0.359). As seen for the dHPC, the mean theta frequency in the range of 7-10 Hz of the mPFC also increased during broadband noise stimuli (8.00 ± 0.56 Hz) compared to during silence (7.23 ± 0.65 Hz, p=0.002, Fig. 7D-E). Power spectrum analysis also revealed a small decrease in the mean power of theta, from mPFC theta power in silence of 430.89 ± 119.18 μV^2^/Hz to 384.78 ± 126μV^2^/Hz during noise stimulation (p=0.0025, Fig. 7F). When analyzing the simultaneous recordings of the mPFC and dHPC, from mice running on the treadmill, showed that dHPC-mPFC theta (7-10Hz) coherence was significantly higher during the noise protocol (0.473 ± 0.152) compared with in silence (0.402 ± 0.140, p=0.030, Fig. 7G-I). This shows that loud noise increased the synchrony in theta (7-10Hz) of dHPC-mPFC during running.

**Figure 7.**
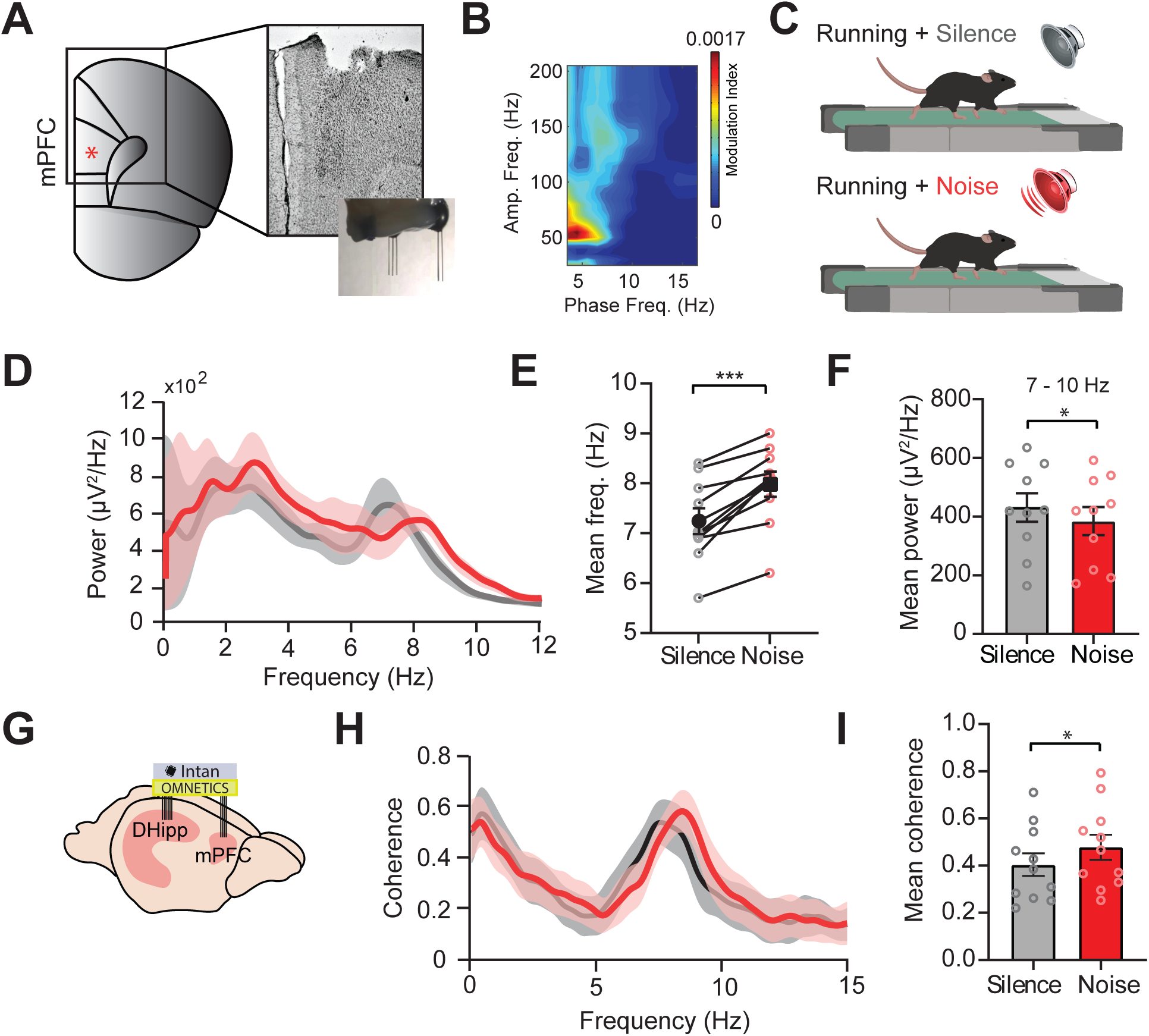
Broadband noise increases type 1 theta (7-10Hz) frequency in the mPFC and coherence between dHPC and mPFC. A) Schematic drawing of the prelimbic mPFC (*red) and brightfield image showing the entry point of the array aiming at the prelimbic area. Lower inset; photo of the dual array for recording the dHPC and mPFC. B) Cross-frequency coupling spectrogram of the LFP signal in the mPFC confirming the expected location of the electrode array. C) Experimental design. D) Mean PSD plot of LFP signal (0-15Hz) recorded in the prelimbic mPFC while mice (n=10) run on a treadmill during silence (gray) and noise stimuli (red). Solid lines show the mean signal, and colored shading shows ±LSEM. E) Line graph showing an increase in mean frequency during noise (red) compared to silence (gray) for slow theta oscillations of the mPFC (p=0.0253, n=10). F) Bar graph showing that mean theta power was significantly different between silence (gray) and noise stimuli (red), p=0.0002, n=10. G) *Left:* Schematic drawing of electrodes for recording in the mPFC and dHPC. *Right:* Mean dHPC/mPFC coherence when running on a treadmill in silence (gray) and during noise stimuli (red). H) Bar graph showing the coherence of 7-10 Hz oscillations during silence (gray) and loud noise recorded in the dHPC and mPFC (p=0.0306). Statistical tests error bars show SEM, *p < 0.05, ***p < 0.001.

## Discussion

The present data provide compelling evidence for a direct, auditory cortex-independent route by which loud noise can rapidly modulate hippocampal activity, with implications for understanding threat detection during movement. Here we show that loud broadband noise accelerates theta (7 - 10 Hz) frequency while decreasing its power during treadmill locomotion. This result aligns with previous studies showing that neurons in limbic regions are activated by high-intensity broadband noise [7–9,48,49]. Our results could not be reproduced when using a tonal stimulus (20 kHz) consistent with previous studies showing that hippocampal responses to tones are weak [8,9,48,49]. For instance, loose-patch experiments revealed that approximately 15% of CA1 neurons respond to noise bursts in awake mice, whereas less than 1% respond to tones [9]. While in CA3, neurons exhibited a stronger response to broadband noise compared to other sound stimuli, suggesting that the auditory information reaching this area is predominantly carried by broadband signals [49]. Studies have shown that hippocampal theta frequency decreases in response to novel environments but increases as the environment becomes more familiar [50,51]. Here we reversed the order of stimulus presentation, introduced noise before silence, or repeated the task over five consecutive days, and found theta frequency variations still persisted independently of novelty/familiarity. Moreover, our findings resonate with a recent study demonstrating that human hippocampal theta frequency increases with higher visual sensory information density during virtual locomotion, a response more pronounced than that of velocity alone [52]. Using a treadmill with a variable speed control, we observed that noise-induced theta acceleration was restricted to the initial two minutes of sound presentation. This period included both the acceleration (0–12 cm/s) and the constant speed (12 cm/s) phase, with no significant changes observed during deceleration. These results suggest that noise effects on theta dynamics may be particularly salient during periods of active locomotor engagement. Still, one limitation of our study is that we used constant noise stimuli, in respect to loudness and duration, yet this stimuli was still enough to consistently accelerate locomotion-induced theta oscillations.

Noise-induced decrease in theta power might reflect desynchronization of the dHPC by the CA1 interneurons and pyramidal cells that directly respond to loud noise [9]. Interestingly, despite the reduced number of noise-responding neurons, the optogenetic stimulation of DCN neurons projecting to the dHPC led to a significant decrease in theta power and an increase in theta frequency. In addition to supporting the hypothesis that auditory stimuli engage brainstem-hippocampal pathways to modulate hippocampal dynamics [7], our results indicate that changes in theta dynamics are not solely dependent on the magnitude of neuronal recruitment, but may instead reflect intrinsic adjustments of the network to external stimuli. For example, reductions in cortical low-frequency LFP power and network desynchronization have been linked to memory formation and retrieval [53], spatial learning [54], associative memory encoding [55] and fear memory [9]. Here we also show a decrease in gamma (40 - 70Hz) power in response to loud noise. This suggests that noise-induced network desynchronization may span multiple frequency bands; however, further investigations are required to elucidate the mechanisms underlying these responses.

### Noise processing in limbic areas compliments the auditory pathway

Our study found that lowering A1 pyramidal cell activity during broadband noise did not remove the noise-driven increase in locomotion-related theta frequency. Our results align with a previous study showing that response latencies to broadband noise in A1 and dHPC are similar, suggesting simultaneous and independent processing between these regions, while the MS and EC showed shorter response latency, highlighting their role in the initial mediation of auditory noise stimuli [9]. Moreover, sound-evoked responses in the auditory cortex are generally weaker during locomotion compared to immobility [3–5,28,56]. For example, optogenetic stimulation of cortical motor neuron axon terminals in the auditory cortex suppressed both spontaneous and stimulus-evoked activity of auditory neurons, providing evidence of a direct cross-modal interaction [5]. In addition, recent findings reveal that neuronal ensembles of the auditory cortex simultaneously encode sound and locomotion speed, enabling behaviorally relevant auditory perception during movement [57]. Navigation-related areas may facilitate the rapid sensorimotor integration of auditory processing by either operating in parallel with the auditory cortex or directly contributing to sensorimotor integration within it. Acoustic motor integration is also apparent in the acoustic startle response, where pontine reticular nucleus neurons are excited by acoustic stimulation of higher intensities (80 dBSPL) via the cochlear nucleus [58,59]. Here we found that optogenetic stimulation of DCN neurons projecting to the PRN [8,60] could mimic the broadband noise response in the dHPC during locomotion. Hence, our results functionally confirm participation of DCN and PRN neurons in the reticular-limbic auditory pathway [8], and that it is possible to induce an electrophysiological response in the dHPC when stimulating an auditory/somatosensory relaying region as the DCN in the absence of sensory stimulation.

### Both the entorhinal cortex and MS PV+ neurons can modulate locomotion-induced theta oscillations in a noisy environment

The dHPC receives sensory input via the EC [61] and plays a critical role in spatial navigation and contextual encoding. Here, we found that when inhibiting excitatory neurons of the MEC and LEC bilaterally, the increase in theta frequency during broadband noise is abolished while the decrease in power persisted, pointing to the role of the EC in integrating multimodal sensory information from the environmental context [12,62]. Along the same lines, it was recently demonstrated that the MEC encodes spatial information from visual and auditory stimuli during locomotion, creating distinct cognitive maps for each sensory modality [63]. Interestingly, we also observed a reduction in theta power during silence when the MEC and LEC were inhibited. Similarly, MEC lesions lead to a decrease of theta oscillation amplitude in the hippocampus [64]. Layer II of the entorhinal cortex (EC-II) is thought to drive theta interactions and synchronization in the hippocampus during novelty exploration, thereby facilitating the integration of external information [65]. Here, we find that chemogenetic lowering of EC activity decreases theta power, but that this modulation is independent of auditory stimuli.

Glutamatergic MS cells are believed to serve as the primary relay for broad-band noise information to the EC via the non-canonical pathway and blocking MS neuronal firing with lidocaine abolishes noise-evoked responses in both the EC and dHPC [8,9]. MS GABAergic and glutamatergic neurons are known to linearly modulate theta oscillation and speed cells of the dHPC and EC [21–23,25,30,66]. However, MS cholinergic neurons infer a complex influence on theta as increasing the MS slow-firing ChAT-positive neurons activity reduces theta frequency in the MEC but have little direct effect on hippocampal theta in awake, active mice and rats [26,67,68]. In our study, inhibition of MS cholinergic neurons did not influence dHPC theta frequency or power in silence nor during broadband noise. As for GABAergic cells, PV+ projection neurons from the MS to dHPC are thought to drive hippocampal theta oscillations [23,69]. Here we demonstrated that the stimulation of MS PV+ neurons increases theta frequency while decreasing its power in the dHPC during silence, mirroring the effect of noise exposure in dHPC theta oscillations. In agreement, rhythmic stimulation of MS PV+ neurons increases theta frequency in freely behaving mice [30]. Furthermore, MS PV+ projection neurons play a crucial role in sensory perception, with septo-hippocampal GABAergic boutons responding to locomotion and sensory stimuli in a manner proportional to stimulus intensity [70].

The MS is known to mediate aversion to auditory and somatosensory stimuli, where glutamatergic neuron ablation decreases aversion, while GABAergic cell ablation enhances it (Zhang et al., 2018). These findings suggest that there is an interaction between MS glutamatergic and PV+ neurons in modulating hippocampal theta response to loud noise, potentially linking sensory processing with emotional responses. Additionally, rhythmic stimulation of MS PV+ neurons was recently shown to directly control various oscillatory frequencies of the MEC [71], and GABAergic projections from the MS selectively inhibit interneurons in the MEC [22]. Furthemore, MS PV+ projection neurons inhibit low-threshold interneurons in the MEC, disrupting theta-rhythmic firing of MEC neurons [28]. However, another study found that only theta power was significantly affected by MS-GABAergic inhibition [27]. Our results suggest that the EC and MS PV+ neurons play a critical and interconnected role in modulating theta dynamics during noise. Furthermore, MEC and LEC separately may provide different mechanisms for modulating theta activity in this context, which should be further investigated.

### Loud noise modulates calcium dynamics of dHCP pyramidal neurons

Our calcium imaging data shows that pyramidal cell activity decreases dramatically (63%) when loud noise animal is running. We demonstrated that running in loud noise decreased calcium activity in 63% of pyramidal neurons in the dHPC, while increasing in 33%, when compared to running in silence. Notably, the small population of loud noise-sensitive neurons aligns with findings by Xiao et al. (2018)[9], who reported that only 15% of CA1 neurons showed heightened activity in response to noise. More so, when mimicking noise modulation of the dHCP with optogenetic activation of DCN neurons, projecting to the PRN, resulted in a decrease in calcium activity in 74% of pyramidal neurons and an increase in 21%, similar to noise response. Additionally, we observed a decrease in the average calcium activity of pyramidal neurons during both noise exposure and optogenetic stimulation of the DCN-PRN pathway. This finding supports the hypothesis that auditory stimuli engage brainstem-hippocampal pathways to modulate hippocampal dynamics [7,8].

The observed reduction in neuronal activity during noise exposure, may reflect increased inhibitory activity and desynchronization of neurons during loud noise. A similar mechanism was described in the auditory cortex (A1), where cortical neurons adapt to background noise by increasing response thresholds and latencies, or shifting rate-level functions toward higher sound levels, thereby enhancing the alignment with the acoustic context [72–76].

A recent study showed that CA1 inherits context-dependent sensory information from the CA3 and EC inputs, to facilitate navigation and memory encoding [77]. Thus, the observed reduction in pairwise correlation during noise in our study may represent circuit-level mechanisms that prevent over-synchronization and preserve flexibility in adapting to the sensory environment [78]. Nonetheless, the low pairwise correlation of pyramidal neurons observed across all analyzed contexts is consistent with previous findings, which report low population synchrony during theta-related exploration and higher synchrony during slow-wave sleep [79].

### Loud noise increase synchrony between dHCP GABAergic interneuron

Noise-driven inputs amplify interneuron connectivity to prioritize behaviorally relevant signals, consistent with findings that GABAergic interneurons coordinate network responses during sensory processing [80,81]. Here we found that the correlation between neurons in Cluster 1, which contained the majority of neurons (44.8% of total), increased significantly during noise compared to silence, potentially reflecting a circuit-wide adjustment to processing auditory stimuli. Furthermore, we found an increase in calcium activity in these neurons specifically during the first minute of loud noise exposure, compared to in silence. Optogenetic activation of hippocampal PV+ interneurons has been shown to broadly inhibit pyramidal neurons while simultaneously enhancing theta frequency resonance in a subset of cells [81]. This mechanism could account for the observed increase in theta frequency during the initial minutes of noise exposure in our data. Additionally, GABAergic neurons in the MS innervates a substantial proportion of hippocampal GABAergic interneurons [82] which supports disinhibition of a subgroup of dHPC pyramidal neuron by auditory stimuli recruiting non-canonical pathways to modulate hippocampal dynamics.

Broadband noise stimulation caused a decrease calcium activity of GABAergic interneurons of the first cluster (Figure 6) after the first minute, and in other clusters throughout the entire stimulus duration. It was demonstrated previously that silencing PV interneurons in the hippocampus did not affect burst firing but shifted the spikes’ theta phase toward the trough of theta [83]. This aligns with our electrophysiological findings, which show an increase in theta frequency despite a decrease in power. Furthermore, sustained photoinhibition of PV+ interneurons or SST+ interneurons has been shown to suppress gamma oscillatory power [84], which could explain the reduction in gamma power observed in our electrophysiology data during noise exposure. The modulation of CA1 pyramidal neurons, as well as distinct interneuron clusters, underscores the role of the hippocampus as a multimodal integrator.

### Increase coherence between the dHPC and mPFC may contribute to alertness

The mPFC, hippocampus, and amygdala are essential regions for anxiety and fear responses [85] to sound stimuli [9,86,87]. Here, we showed that broadband noise stimuli also increased mPFC fast theta frequency and decreased its power. A recent study revealed that the mPFC contains neurons that respond to sound, especially high-intensity broadband noise during wakefulness [87]. Moreover, here we show that broadband noise induces an increase in the type 1 theta coherence between mPFC and dHPC. These areas are highly interconnected, with multiple reciprocal projections facilitating interactions of anxiety and fear circuits [88], and it has been shown that dHPC–mPFC theta coherence reflects novelty-induced selective attention and short-term habituation [89]. Our results suggest that the response of type 1 theta in the mPFC to broadband noise is derived from the dHPC and may reflect the relative novelty of the noise stimuli in the environment, contributing to attention and an alert state. The enhanced coherence between the dHPC and mPFC suggests that noise strengthens interregional synchrony, potentially prioritizing processing of behaviorally relevant stimuli.

## Conclusion

This study highlights the hippocampus as a dynamic hub that integrates auditory stimuli and modulates neural activity based on context and behavioral demands. In summary, these results expanded our understanding of how the reticular-limbic pathway can integrate loud noise into neuronal processing of the hippocampus, and we speculate that the functional output of increased theta 1 frequency during running is to make animals more alert in response to possible hazards.

## Material and Methods

### Animals

Six to seven weeks old male C57BL/6 mice (n= 67), Choline acetyltransferase (ChAT)-Cre mice (n=5, ChAT-IRES-Cre; stock #006410, Jackson Laboratories), and parvalbumin (PV)-cre (n=4, PV-IRES-Cre; stock #017320, Jackson Laboratories) were housed on a 12h/12h day/night cycle to maintain their normal biorhythms and had free access to food and water. All procedures were approved and followed the guidelines of the Animal Ethics Committee (CEUA) of the Federal University of Rio Grande do Norte (protocol numbers 052/2015 and 135.064-2018) and Uppsala University (protocol number 5.8.18-08464/2018). Efforts were made to minimize the suffering and discomfort of animals and to reduce the number of animals used. Experiments were carried out during the dark phase (∼19:00 - 22:00 pm).

### Virus injections

Approximately four weeks before chemogenetics, optogenetics, or calcium imaging experiments, mice were injected with viral constructs using chemogenetic viral vectors for expressing mutated muscarinic M4 receptors (hM4Di), optogenetic viral vectors for expressing opsins coupled to a channelrhodopsin2 (ChR2), or genetically encoded calcium indicator (GCaMP6f), respectively. In detail, mice were anesthetized with isoflurane (3% for induction and 1% for maintenance) and placed on a 37°C heating pad controller. The skin was anesthetized with lidocaine hydrochloride 3% before a straight incision was made and a small hole was drilled with a dental micro drill (Beavers Dental) using the coordinates of the region of interest, detailed below. Next, aliquoted virus was withdrawn into a microsyringe (10Lμl Nanofil, 34-gauge removable needle) at the speed of 1.5Lμl/min using an infusion pump (Chemyx NanoJet). The syringe was lowered into the area to the deepest dorsoventral (DV) coordinate (for vertical extending areas), and 0.75Lμl of virus of interest was slowly infused (1.5Lμl/min) and the needle was kept in place for 10 min before retracted to the second, more superficial DV coordinate, for a second infusion (0.75Lμl) of virus. After 10 min, the syringe was carefully withdrawn and the skin was sutured, with additional lidocaine hydrochloride 3% applied over the suture.

For experiments using chemogenetic inhibition viral vectors of AAV9.CaMKIIL.hM4D(Gi).mcherry (Addgene,100 µL at titer ≥1×10¹³ vg/mL, dilution 1:3 in saline) were bilaterally injected into the auditory cortex of C57BL/6 mice (0.75 μl per injection site) at two different anteroposterior coordinates (AP: -2.75 and -3.1 mm, ML: ±4.0 mm, DV: -2.0 mm) to cover a large area of the primary auditory cortex. For chemogenetic inhibition of the entorhinal cortex both the lateral entorhinal cortex (LEC) and medial entorhinal cortex (MEC) were bilaterally injected (0.75 μl per hemisphere) at the coordinates for the MEC; AP: -4.70 mm, ML: ±3.33 mm, DV: -3.3 mm [67] and for the LEC; AP: -3.2 mm, ML: 4.6 mm, DV: -3.6 mm [90].

For optogenetic stimulation of the dorsal cochlear nucleus (DCN) neurons projecting through the pontine reticular nucleus (non-canonical pathway), an AAV retrograde vector (AAVrg-CAG-hChR2-H134R-tdTomato, Addgene, 100µL at titer ≥7×10¹² vg/mL, dilution 1:3 in saline) was injected (0.75 μl per hemisphere) into the pontine reticular nucleus of C57BL/6 mice at the following coordinates: AP: -5.2 mm, ML: ±0.7 mm, DV: -4.0 and -4.5 mm [8], to be able to excite channelrhodopsin 2 (ChR2)-expressing DCN neurons with blue light to excite the DCN – pontine reticular nucleus pathway.

For examining the contribution of different neuronal populations of the medial septum in modulating response to loud broadband noise, two different viral constructs were injected into two different transgenic lines at the coordinates AP: +0.7 mm, ML: 0.5 mm, and in a 12° angle towards the midline at two different depths of DV: -3.5 and -3.8 mm [67]. For chemogenetic inhibition of MS cholinergic neurons, ChAT-cre mice were injected with AAV9.Syn.DIO.hM4D(Gi).mCherry (Addgene, 100 µL at titer ≥1×10¹³vg/mL, dilution 1:3 in saline). For optogenetic activation of GABAergic MS neurons, PV-cre mice were injected with pAAV9-EF1a-double-floxed-hChR2(H134R)-mCherry-WPRE-HGHp (Addgene, 100 µL at titer ≥1×10¹³vg/mL, dilution 1:3 in saline).

For dHPC calcium imaging, mice were injected with genetically encoded calcium indicators with either the calcium/calmodulin-dependent protein kinase II (CaMKII) or synapsin promoter (pENN.AAV.CaMKII.GCaMP6f.WPRE.SV40, Addgene, titer ≥1×10¹³vg/mL or pAAV.Syn.GCaMP6f.WPRE.SV40, Addgene, titer ≥1×10¹² vg/mL) to target CA1 hippocampal pyramidal cells (PCs) at the following two coordinates: AP -1.65 mm, ML +1.85 mm, DV 1.2 mm; and, AP -1.8 mm, ML +1.7 mm, DV -1.3 mm (0.75 µL/site). To target hippocampal CA1 interneurons, viral constructs containing the mDlx promoter region were used (AAV9.mDlx.GCaMP6f, titer ≥ 1×10¹² vg/mL).

### Chemogenetic pharmacology

Neurons expressing the G_i_-protein coupled mutated muscarinic receptors 4 (hM4Di) were activated by low-dose (0.5 mg/kg i.p; [91]) clozapine-N-oxide (CNO) and reduced neuronal activity was examined 40 min after the injection. Local expression of CaMKIIL.hM4D(Gi).mcherry was routinely confirmed post hoc by slicing brains in 50μm, sections and examined by fluorescence microscopy.

### Chronic electrode, optic fiber and miniscope lens implants

Two types of electrode arrays were fabricated from insulated tungsten wires (30 µm, impedance between 300-800 kΩ, California Wires). For the dHPC a 6-channel electrode (2×3, electrode spacing 200 μm) was used, while for the dual targeting of the dHPC and the prelimbic region of the mPFC a 10-wire array was custom-made so that 6 wires (2×3, 200 μm spacing) would target the dHPC, and a 4 wire electrode (2×2 μm, 200 μm spacing) was used for targeting the mPFC. Mice were implanted with electrodes as previously described[92,93]. Coordinates for the dHPC array were AP: 1.8 mm, ML: 1.7 mm and DV: 1.2 mm. For the dual area targeting electrode, the coordinates for targeting the mPFC were AP: 2.1 mm, ML: 0.3 mm and DV: 1.6 mm. A micro screw placed into the skull covering the cerebellum served as the reference electrode.

Optic fiber implants were fabricated from 200 um diameter fiber (Thorlabs, USA), and ceramic ferrules had a diameter of 230 um (Thorlabs, USA). For light stimulation of medial septum, an optical fiber was implanted at the coordinates AP: +0.8 mm, ML: ±0.3 mm, and DV: 3.5 mm) at a 10° angle. For optogenetic stimulation of DCN neurons projecting to the pontine reticular nuclei, optical fibers were bilaterally implanted above the DCN (AP: - 6.1 mm, ML: ±2.3 mm, and DV: -3.9 mm). Fibers were implanted during the same surgery as for electrode arrays. Following surgery, animals were housed individually and allowed to recover for at least 10 days before further experiments.

For dHPC calcium imaging, ten days after the virus injection, mice underwent surgery for implanting the gradient index (GRIN) lens, as previously described in detail (Winne et al., 2021). In brief, the cortex was aspirated above the right hippocampus area until the white fibers of the corpus callosum were visualized (as the working distance of the miniscope is 0.3 to 0.5 mm, the lens can be placed on the corpus callosum). The GRIN lens (length: 5.0 mm; diameter: 2.0 mm; pitch: 0.25; object working distance: 0.2 mm, Edmund Optics Corporation) was implanted at the coordinates AP: -2.70 mm, ML: +3.95 mm, and DV: 2.70 mm. Two weeks later, GCaMP6f expression was confirmed through the lens in anesthetized mice fitted in the stereotaxic frame using the miniscope. Next, a base plate is magnetically attached to the miniscope and lowered over the lens, and the optimal image plane is identified. Next, the baseplate is fixed to the skull using UV-hardened cyanoacrylic glue (Winne et al., 2021), and the miniscope is gently removed. Following surgery, animals were housed individually and allowed to recover for at least 10 days before further experiments.

### Treadmill locomotion

We used an electrical treadmill (9 × 26.6 cm, Columbus Instruments) with controlled speed. Individual animals were habituated to the equipment for 2 consecutive days where animals were allowed to freely explore the treadmill turned off for 5 min and next trained on the treadmill to perform voluntary 3 min runs, with the speed gradually increasing from 0 to 12 cm/s (first minute), kept constant at 12 cm/s (during 1 min) and decreasing from 12 to 0 cm/s (last minute). We also used a shorter 140-second protocol just with the initial group (Figure1, D-H), during which the treadmill speed was gradually increased from 0 until it reached 12 cm/s. At the beginning of each recording session, animals were connected to a headstage and placed individually at the center of the treadmill.

### Acoustic and light stimulation

Acoustic Stimulation during the locomotion consisted of broadband noise (1–20 kHz ± 0.5 kHz, 80 db sound pressure level, during 180 s) or a pure tone (20 kHz ± 0.5kHz, 90 db, 180s in duration). Sounds were created using a custom script in Matlab (version 2020a, MathWorks). The synthesized signals were amplified using a (Taramps TS400×4, Brazil) and delivered through a loudspeaker (Selenium Trio ST400, JBL by 71 Harman, Brazil) placed 15 cm above the mouse. All experiments were carried out in a sound shielded behavior room. In an attempt to mimic broadband noise, the optogenetic light stimulation (blue light, using a 473 nm emission laser) was generated as a waveform with random components (frequency and amplitude) during 180 s.

### Electrophysiology

Local field potentials (LFPs) were acquired using a 16-channel amplifier (Intantech) and custom software modified from the Intan RHA evaluation package (Intan technologies) or the OpenEphys[94] software with an Intan RHD headstage. The Intan RHA headstage has a fixed 1 Hz high pass filter, while the RHD headstage has a high pass filter of 0.1 Hz. Differences in filtering settings did not affect low-frequency oscillatory power (2 -10 Hz) as evaluated by synthetic signals. During LFP recordings, mice walking on the linear treadmill were simultaneously video recorded using a Basler camera (model acA1300-30um), with the data acquisition system also collecting digital trigger pulses. A maximum of three mice were recorded per day.

### Calcium imaging

At the beginning of each imaging session, the animals were briefly anesthetized with isoflurane (3%) to fit the miniature microendoscope (Miniscope V3, UCLA) [95] to the magnetic base plate. The imaging session started 20 min after the mice recovered from anesthesia. The focus of the miniscope was manually adjusted to an imaging field ∼200μm away from the lens. Optimum laser power (20% of the maximum), imaging gain, and focal distance were selected for each animal and conserved across all sessions. The Ca^2+^ signal was acquired at a frame rate of 30Lframes per second using the UCLA V3 Miniscope controller software. Two consecutive days before the first recording session, the mice were habituated to wearing the miniscope, experimental room, and walking freely in the treadmill apparatus for 10 minutes.

### Electrophysiology analysis

The theta power was correlated with locomotion speed by downsampling total theta power vs. time obtained from a spectrogram (Matlab spectrogram command) of a recording channel placed at the *stratum radiatum* of the dorsal hippocampus. Power spectral densities (PSD) for all channels were computed using the Welch method (pwelch Matlab command). In the experiments recording from both the mPFC and dorsal hippocampus the coherence between these areas was calculated using the Matlab command Mscohere. All custom software can be found at http://github.com/cineguerrilha/Neurodynamics.

### Calcium imaging post-processing and analysis

Calcium activity was extracted using Minian [96], an open-source analysis pipeline for single-photon calcium imaging data based on the CNMF algorithm [97,98]. Miniscope videos were motion-corrected using NormCorre [99], spatially downsampled by a factor of 2, and temporally downsampled by a factor of 2 to reduce noise and the computational power required for cell segmentation. Spatial and temporal components for individual cells were extracted using large-scale sparse non-negative matrix factorization (CNMF/CaImAn, [98]). To find the same cells in different experimental sessions recorded on the same day (for example, silence vs. noise), the videos from all sessions of the same animal were preprocessed together in the Minian Pipeline.

Analysis of calcium dynamics from individual neurons (filtered calcium traces) was conducted using custom scripts in Matlab (Mathworks). Calcium activity is reported as the change in fluorescence intensity relative to resting fluorescence intensity (ΔF/F), and it was normalized for each animal. The altered or unchanged cellular activity criteria combine statistical significance and percentage change. Neurons whose changes in activity during noise or optogenetic stimulation showed a p-value < 0.05 in the t-test are considered statistically significant. Those with a percentage change greater than 5% are classified as having increased activity, and those less than -5% as having decreased activity. Neurons whose percentage changes are between -5% and 5% or that do not reach statistical significance are classified as having no significant change. The neuronal activity traces were organized according to the K-means clustering solution for clustering. The K-means clustering algorithm was applied, and the number of predefined clusters based on the silhouette analysis was used to evaluate the clustering quality. Neurons with silhouette values below 0.1 were excluded from the final analysis.

### Statistical analysis

For electrophysiology, the normality of the data was tested using the Shapiro-Wilk test and the sphericity with the Mauchly’s test, with additional Geisser-Greenhouse correction if the sphericity was violated. We used One-way repeated measures ANOVA with Tukey’s HSD test for multiple comparisons to calculate the statistical significance of normally distributed data. For non-normal distributed data, we used the nonparametric Wilcoxon matched pairs or Friedman’s test to calculate statistical significance [100]. For calcium imaging the statistical significance of normally distributed data, we used the paired T-test. Statistical significance was defined with α < 0.05. Data is presented as mean ± standard error of the mean (SEM).

## Funding Sources

American Tinnitus Association (ATA), Brazilian Research Council, Coordination for the Improvement of Higher Education Personnel (CAPES) and the Brazilian National Council for Scientific and Technological Development (CNPq).

## Conflict of interest

The authors declare no conflict of interest.

## Author contribution

JW and RNL designed the study; JW and MN did experiments; JW and RP analyzed the data; GN and RNL constructed the custom hardware; JW and CSS designed figures; JW and KEL wrote the manuscript with input from CSS, KK and RNL.

## Notes

### Competing Interest Statement

The authors have declared no competing interest.

