## Supplementary figures and images for "Auditory regulation of hippocampal locomotion circuits by a non-canonical reticular-limbic pathway"

### Supplementary Fig. 1

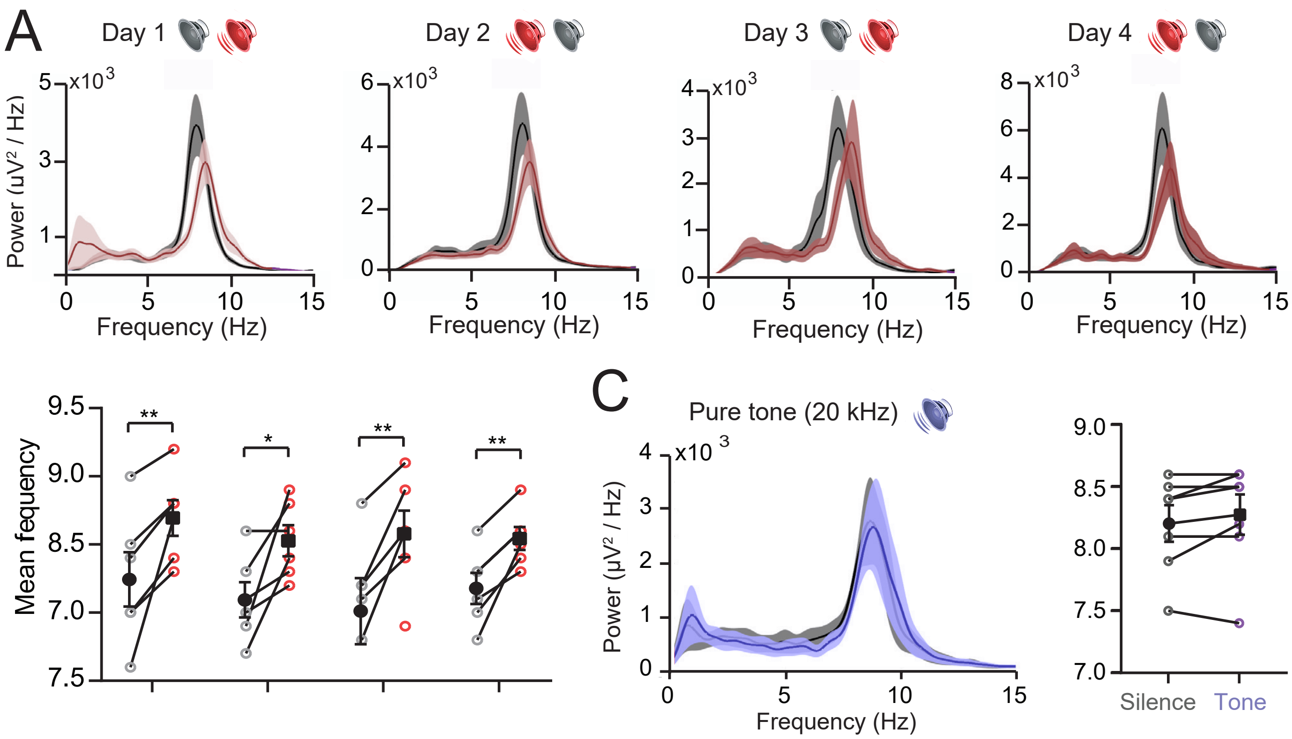


Supplementary Figure 1
